# A phylogenetic and evolutionary analysis of antimycin biosynthesis

**DOI:** 10.1101/164145

**Authors:** Rebecca Joynt, Ryan F. Seipke

## Abstract

*Streptomyces* species and other *Actinobacteria* are ubiquitous in diverse environments worldwide and are the source of, or inspiration for, the majority of pharmaceuticals. The genomic era has enhanced biosynthetic understanding of these valuable chemical entities and has also provided a window into the diversity and distribution of natural product biosynthetic gene clusters. Antimycin is an inhibitor of mitochondrial cytochrome c reductase and more recently was shown to inhibit Bcl-2/Bcl-X_L_srelated anti-apoptotic proteins commonly overproduced by cancerous cells. Here we identify 65 putative antimycin biosynthetic gene clusters (BGCs) in publicly available genome sequences of *Actinobacteria* and classify them based on the presence or absence of cluster-situated genes *antP* and *antQ,* which encode a kynureninase and phosphopantetheinyl transferase (PPTase), respectively. The majority of BGCs possess either both *antP* and *antQ* (L-form) or neither (S-form), while a minority of them lack either *antP* or *antQ* (I_Q_ or I_P_form, respectively). We also evaluate the biogeographical distribution and phylogenetic relationships of antimycin producers and BGCs. We show that antimycin BGCs occur on five of the seven continents and are frequently isolated from plants and other higher organisms. We also provide evidence for two distinct phylogenetic clades of antimycin producers and gene clusters, which delineate S-form from L- and I-form BGCs. Finally, our findings suggest that the ancestral antimycin producer harboured an L-form gene cluster which was primarily propagated by vertical transmission and subsequently diversified into S-, I_Q_ and I_P_form biosynthetic pathways.

## Introduction

Microbial natural products, particularly those produced by filamentous *Actinobacteria*, have been a cornerstone of the pharmaceutical industry for more than half a century [1]. The genes encoding natural product biosynthesis are typically grouped together into a gene cluster, which possibly enhances their transmissibility and the evolution of chemical diversity [2]. Little is understood about the forces driving these processes, but access to large datasets of genome sequences provides an opportunity for exploration.

Antimycin-type depsipeptides are a large and diverse family of natural products widely produced by *Streptomyces* species [3]. The family’s namesake, the nine-membered ringed antimycins, were discovered more than 65 years ago [4]. Ring-extended members of this family have also been identified and include: JBIR-06 (12-membered ring), neoantimycin (15-membered ring) and respirantin (18-membered ring) [5–7]. All of these compounds possess antifungal, insecticidal and nematocidal activity, as a result of their ability to inhibit mitochondrial cytochrome c reductase via a conserved 3-formamidosalicylate moiety [8]. Antimycins are used commercially as a fish biocide, but were recently found to be potent and selective inhibitors of the mitochondrial Bcl_2_/Bcl-x_L_related anti-apoptotic proteins, which are over-produced by cancer cells and confer resistance to apoptotic chemotherapeutic agents [9]. To date, the biosynthesis of antimycins has been reported for a myriad of environmental isolates, but it was not until recently that the hybrid non-ribosomal peptide synthetase (NRPS) / polyketide synthase (PKS) biosynthetic pathway that directs their assembly was revealed in a strain of *Streptomyces albus* [10].

The ~ 25 kb antimycin biosynthetic gene cluster (BGC) harboured by *S. albus* is composed of 15 genes organised into four polycistronic operons *antAB, antCDE, antFG* and *antHIJKLMNO* (Fig. 1) [11]. The biosynthetic gene cluster was recently used as the basis for the reconstitution of antimycin biosynthesis *in vitro* [12, 13] and heterologous production using *Escherichia coli* [14] and *S. coelicolor* [15]. The biosynthesis and activation of the unusual starter unit, 3-formamidosalicylate is specified by the genes *antFGHIJKLNO* [12, 14, 16]. The di-modular NRPS, AntC and the unimodular PKS, AntD comprise the NRPS-PKS assembly line, while AntE and AntM are crotonyl-CoA carboxylase/reductase and discrete ketoreductase enzymes, respectively, and AntB is an acyltransferase responsible for the acyloxyl moiety and the chemical diversity observed at R^1^ (Fig. 1) [13]. The expression of the antimycin BGC is coordinately regulated with the candicidin BGC by a LuxR-family regulator, FscRI, which activates expression of *antABCDE* [15]. The *antA* gene encodes a cluster-situated extracytoplasmic function RNA polymerase sigma (σ) factor named σ ^AntA^, which activates transcription of the *antGF* and *antHIJKLMNO* operons [11].

**Fig. 1.**
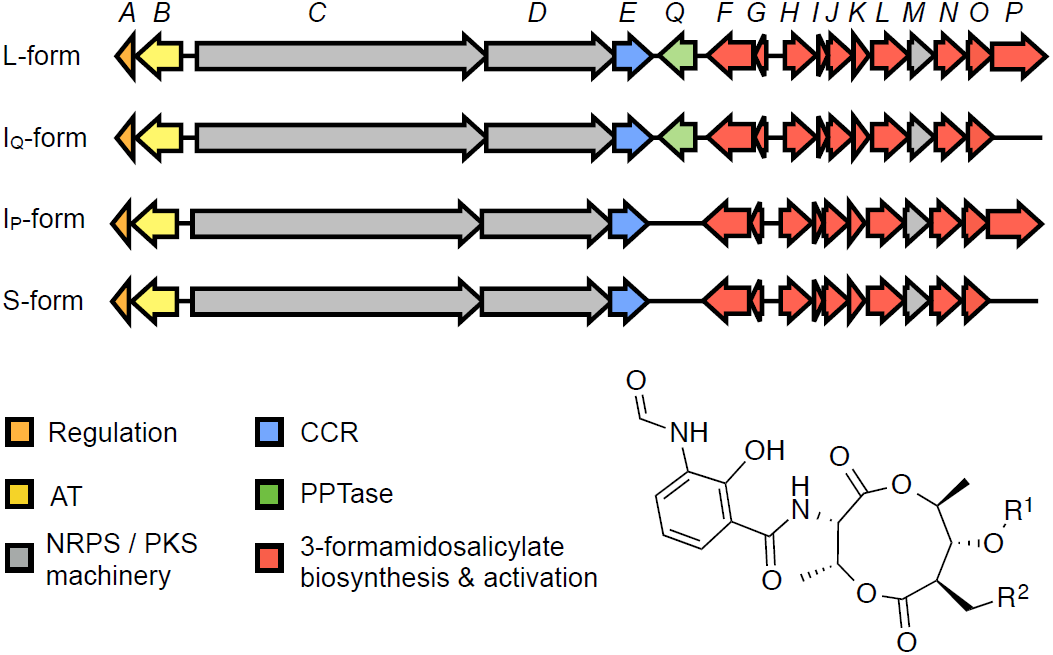
Schematic representation of L-I_Q_ I_P_ and S-form antimycin biosynthetic gene clusters. AT, acyltransferase; NRPS, non-ribosomal peptide synthetase; PKS, polyketide synthase; CCR, crotonyl-CoA carboxylase/reductase; PPTase, phosphopantetheinyl transferase. Antimycins: Antimycin A_1_, R^1^ = COCH(CH_3_)CH_2_CH_3_, R^2^ = (CH_2_)_4_CH_3_; Antimycin A_2_, R^1^ = COCH(CH_3_)_2_, R^2^ = (CH_2_)_4_CH_3_; Antimycin A_3_, R^1^ = COCH_2_CH(CH_3_)_2_, R^2^ = (CH_2_)_2_CH_3_; Antimycin A_4_, R^1^ = COCH(CH_3_)_2_, R^2^ = (CH_2_)_2_CH_3_.

Intriguingly, subsequent identification of antimycin biosynthetic pathways in other taxa revealed that the BGC possesses up to four architectures [17]. Short-form (S-form, 15 genes), intermediate-form (I_Q_ or I_P_form, 16 genes) and long-form (L-form, 17 genes), based on the absence (S-form) or presence (L-form) of two cluster-situated genes, *antP* and *antQ*, which encode a kynureninase and phosphopantetheinyl transferase, respectively. I-form BGCs harbour either *antP* (I_P_) or *antQ* (I_Q_), but not both (Fig. 1). How the antimycin BGC evolved into these various architectures is an intriguing question and one that we sought to address with this study.

Here we identify 73 antimycin BGCs (six known and 67 putative) in publicly available genome sequences of *Actinobacteria* and evaluate their biogeographical distribution and phylogenetic relationships. Isolation metadata suggests that the antimycin BGC has a large biogeographical range, with isolation of putative antimycin producers from at least five continents. Our phylogenetic analyses support the existence of two distinct clades of antimycin producers and BGCs which delineate S-form from L- and I-form BGCs. Finally, our findings suggest that the ancestral antimycin producer harboured an L-form BGC that was primarily propagated by vertical transmission and subsequently diversified into S-, I_Q_ and I_P_form biosynthetic pathways.

## Results and discussion

### Identification of putative antimycin biosynthetic gene clusters (BGCs) in *Actinobacteria*

Established and putative antimycin BGCs were previously identified within the genomes of 14 *Streptomyces* species [17], however casual analyses of the genome sequences available in GenBank suggested that this number is likely to be far greater. In order to formally assess this possibility, 1,421 publically available genome sequences for select *Actinobacteria* genera (i.e. those with a history of natural products production: (*Actinobactera, Actinomadura, Actinospica, Amycolatopsis, Kitasatospora, Micromonospora, Nocardia, Saccharopolyspora, Planomonospora, Pseudonocardia, Salinispora, Streptacidiphilus,* and *Streptomyces*) were downloaded and annotated using Prokka 1.12 [18]. The Prokka annotation enabled the construction of a customised mutligeneblast database, which was subsequently used in conjunction with the *antFGHIJKLMNOPQ* genes from *S. ambofaciens* and multigeneblast 1.1.13 [19] to generate a list of taxa harbouring a putative antimycin BGC. The genes *antFGHIJKLMNOPQ* were selected on the basis that they are essential for antimycin biosynthesis and are conserved in all established antimycin BGCs; *antPQ* were included in order to permit the tentative classification of gene cluster architecture (see below). Close inspection of gene clusters from the candidate list resulted in the identification of an antimycin BGC in 73 taxa (six known and 67 putative) (Table 1). Among these, five are described as non-*Streptomyces* species: *Saccharopolyspora flava* DSM 44771, *Streptacidiphilus albus* JL83, *Streptacidiphilus albus* NBRC 100918, *Actinospica acidiphila* NRRL B-24431 and *Actinobacteria bacterium* OV320 (Table 1).

**Table 1.**
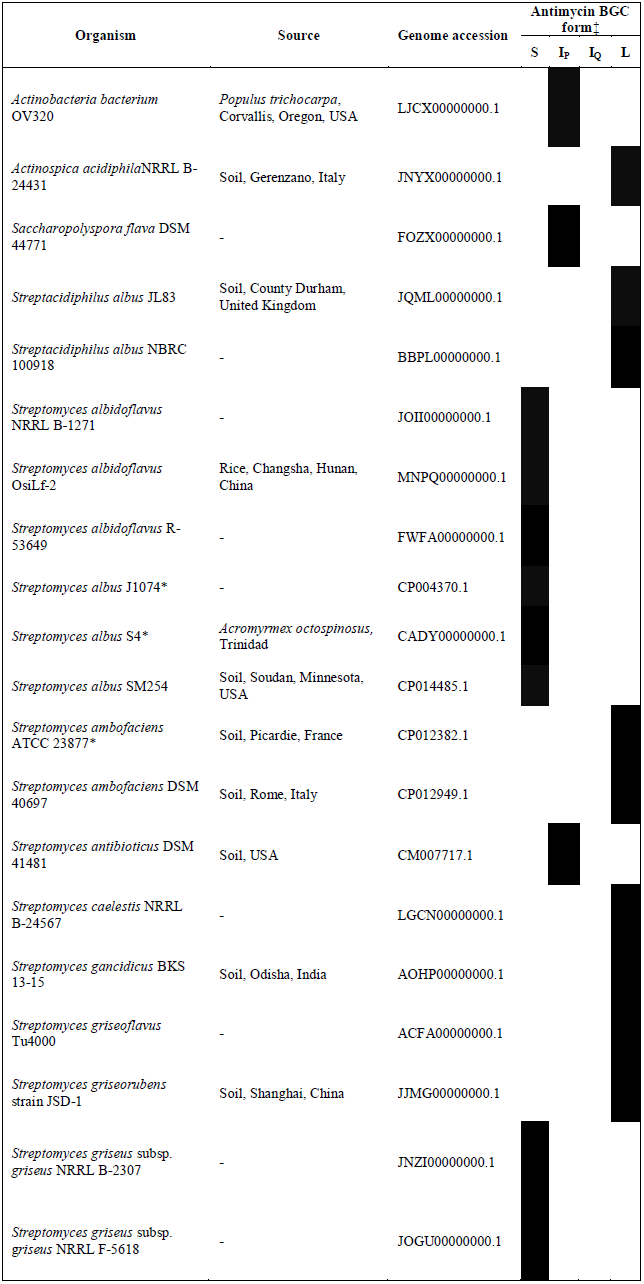
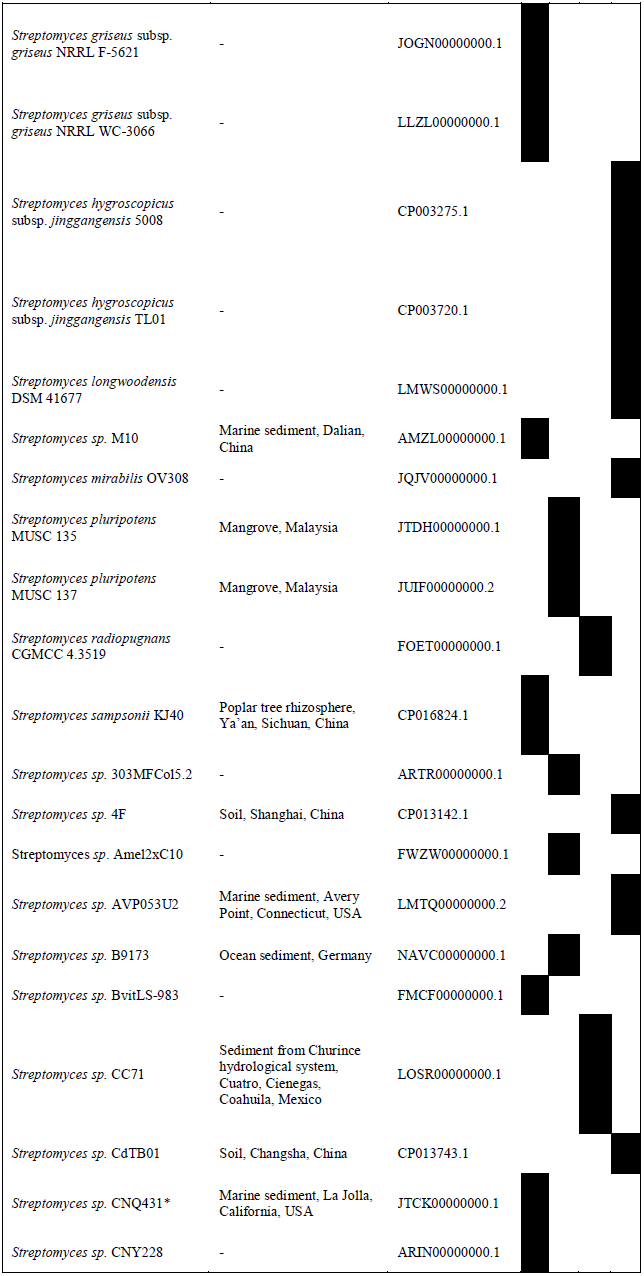
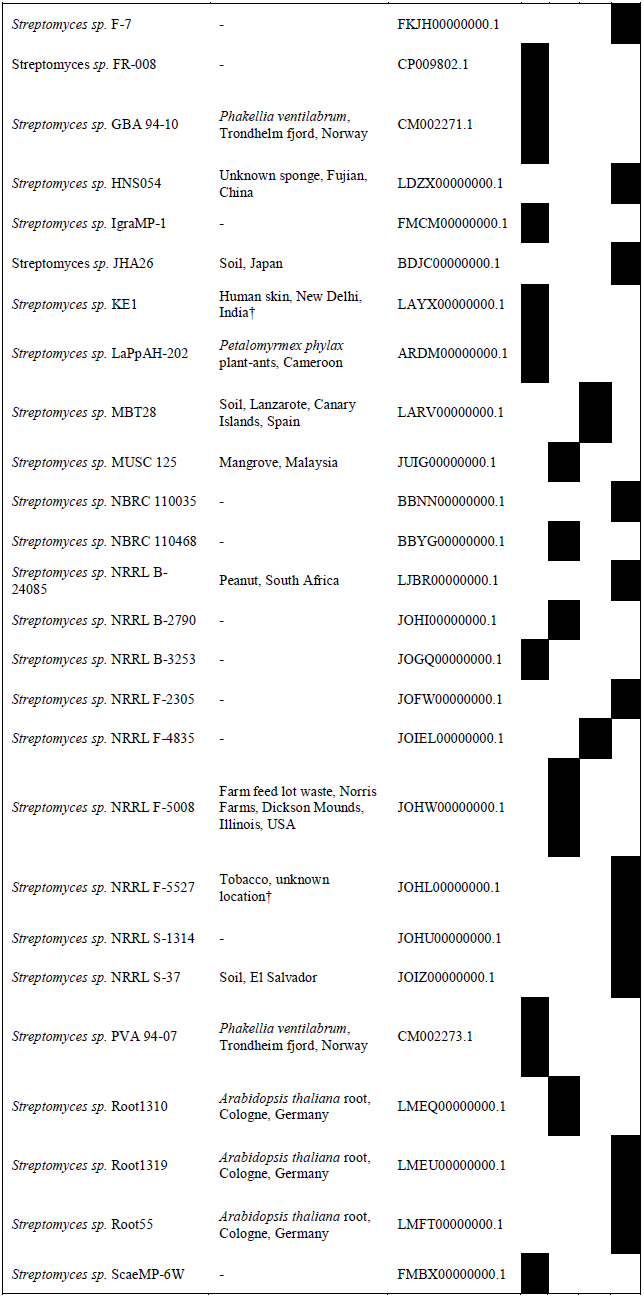
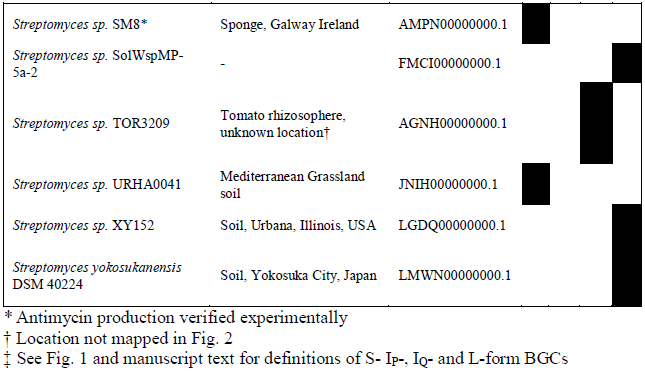
*Actinobacteria* harboring a putative antimycin biosynthetic gene cluster (BGC)

Inspection of loci identified as encoding 3-formamidosalicylate biosynthetic genes revealed a few noteworthy peculiarities. *Streptomyces albus* subsp. *albus* strain NRRL B-2513 possesses a clear 3-formamidosalicylate locus, but lacks *antM* and the core NRPS-PKS biosynthetic machinery at this locus or elsewhere in the genome, and *Streptomyces phaeoluteigriseus* strain DSM 41896 has endured at least two frameshift mutations in *antD,* which presumably render it non-functional. In addition, *Streptomyces lincolnensis* strain NRRL 2936, *Streptomyces* sp. yr375 and *Streptomyces* sp. ERV7 each harbour the same antimycin-like BGC, however gene rearrangement and insertion is evident, for example a small locus of fatty acid anabolism genes has been inserted between the 3-formamidosalicylate biosynthetic genes and the NRPS-PKS machinery, suggesting that the biosynthetic pathway may not in fact produce antimycins. These taxa and BGCs were discarded as a consequence of these peculiarities.

### Classification of antimycin BGCs

Antimycin BGCs exist in four architectures and the gene clusters identified here were classified as short-form (S-form, 15 genes), intermediate-form (I_P_ or I_Q_ form, 16 genes) and long-form (L-form, 17 genes), based on the absence (S-form) or presence (L-form) of two cluster-situated genes, *antP* and *antQ*, which encode a kynureninase (InterPro ID, IPR010111) and a phosphopantetheinyl transferase (InterPro ID, IPR0082788), respectively. The organisation of the genes within antimycin BGCs and their functions are described in Fig. 1 and Table 2, respectively. Annotation of the putative biosynthetic pathways identified above resulted in the classification of 25 S-form, 13 I_P_form, five I_Q_form and 30 L-form antimycin BGCs (Table 1).

**Table 2.** Functions of proteins encoded by antimycin BGCsL-form

| Protein | Function |
| --- | --- |
| AntA | Extra cytoplasmic function RNA polymerase sigma factor |
| AntB | Acyltransferase |
| AntC | Dimodular non-ribosomal peptide synthetase |
| AntD | Unimodular polyketide synthase |
| AntE | Crotonyl-CoA reductase/decarboxylase |
| AntF | Acyl-CoA ligase |
| AntG | Peptidyl carrier protein |
| AntH | Multicomponent oxygenase |
| AntI | Multicomponent oxygenase |
| AntJ | Multicomponent oxygenase |
| AntK | Multicomponent oxygenase |
| AntL | Multicomponent oxygenase |
| AntM | Ketoreductase |
| AntN | Tryptophan 2,3-dioxygenase |
| AntO | *N*-formylase |
| AntP | Kynureninase |
| AntQ | Phosphopantetheinyl transferase |

Interestingly, the conditional loss of *antP* and *antQ* relates to consumption of tryptophan and maturation of the biosynthetic assembly line to its pantetheinylated form. AntF activates anthranilate as the first step in the biosynthesis of the 3-formamidosalicylate starter unit. All forms of the antimycin biosynthetic pathway are able to access anthranilate from the‘core’ anthranilate pool within the cell, evidenced by feeding studies with exogenous fluoroanthranilates [16]. However, only L- and I_P_form pathways harbour a gene encoding a putative kynureninase (AntP), which presumably processes the kynurenine produced by the AntN tryptophan 2,3-dioxygenase to anthranilate. S- and I_Q_-form pathways lack a kynureninase and must rely upon the housekeeping kynureninase involved in tryptophan catabolism. The maintenance of *antP* by L- and I_P_forms and its loss by S- and I_Q_forms may be driven by differences in species-level physiology, for instance the availability of cytosolic anthranilate. It is perhaps not surprising that AntQ is not essential in S- and I_P_forms; in fact, most NRPS and PKS biosynthetic systems lack a cluster-situated PPTase and are dependent on the promiscuity of one or more PPTase enzymes encoded elsewhere in the genome. This is clearly the case for S- and I_P_form antimycin BGCs, but it may not be for L-form antimycin BGCs, as antimycin production by *S. ambofaciens* requires both *antQ* and a PPTase encoded elsewhere in the genome that is similar to SCO6673 from *S. coelicolor* A3(2) [20]. The contextual requirement of *antP* and *antQ* for antimycin biosynthesis creates the opportunity for divergent evolution of the antimycin BGC.

### Biogeographical distribution of antimycin biosynthesis

The biogeography of natural products biosynthesis is an emerging area and one that can not only guide future natural products bioprospecting campaigns, but which enables formulation of interesting questions in chemical microbial ecology [21, 22]. Thus, we curated isolation metadata for putative antimycin producers to ascertain any patterns in source material or its geographical origin. The breadth of data varied considerably, but a source and/or country location was available in GenBank or within the literature for 38 out of 73 strains (Table 1). Sample collection data were plotted onto a World map and pins were colour coded based on gene cluster architecture. Inspection of the resulting map did not show an obvious link between gene cluster architecture and geographical location, but did reveal that putative antimycin producers have been isolated from a relatively large geographical area, including at least five of the seven continents: Africa, Asia, Europe, North America and South America (Fig. 2). Only a single strain originates from the Southern Hemisphere, which is surprising however this is likely a consequence of the inherent limitations of the dataset. Like geographical location, gene cluster architecture and isolation source material do not appear to be related, however, as anticipated, many strains originate from various soil ecosystems (17 in total) or marine sediments (four in total), which supports the longstanding view that these niches are rich sources of bioactive metabolites. Interestingly, 18 of the strains were isolated from plants, sponges or insects, suggesting that they may be involved in symbioses, which is in line with the increasing number of reports implicating antibiotic-producing taxa as defensive symbionts of higher organisms [23, 24]. Overall, these data suggest that antimycin-producing *Actinobacteria* are likely distributed worldwide, which may reflect the significance of producing an inhibitor of eukaryotic cytochrome c reductase in diverse niches.

**Fig. 2.**
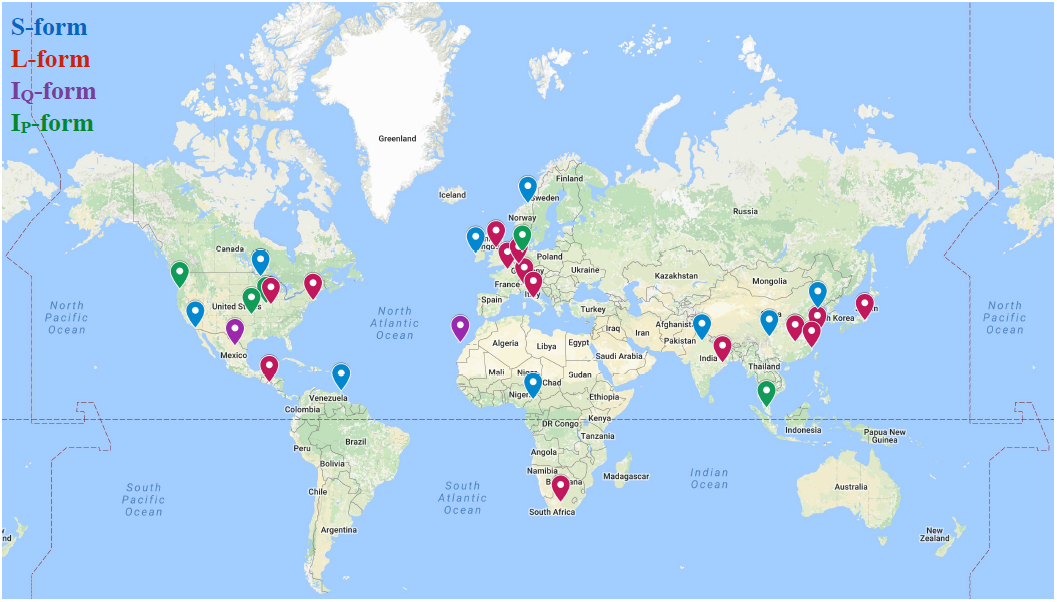
Geographical distribution of 38 *Actinobacteria* harbouring a putative antimycin biosynthetic gene cluster. Map pins are colour-coded based on gene cluster classification. The map is available here: <u>https://drive.google.com/open?id=1rXUFJXSt8szUYMuCJExQbEP4X1k</u>.

### Distribution of antimycin BGCs within *Actinobacteria*

The collection of putative antimycin BGCs identified here provides an opportunity to further explore their phylogenetic and evolutionary relationships. A multilocus phylogeny was reconstructed in order to evaluate the taxonomic distribution of antimycin BGCs. Phylosift was used to identify and extract phylogenetic markers from the genome of each microbe described in Table 1. This resulted in the identification of 29 phylogenetic markers present in single copy in each taxon (see Table S1 for description of markers). The markers were concatenated, aligned (length 13,119 nt) and used to infer a maximum likelihood (ML) phylogenetic tree (Fig. 3). Inspection of the resulting phylogeny suggested that six taxa have been taxonomically miss assigned. For instance, *Actinospica acidiphila* strain NRRL B-24431 and *Actinobacteria bacterium* strain OV320 group closely with *Streptomyces* species placed within the interior of the tree and are therefore likely to be members of the genus *Streptomyces* (Fig. 3). Additionally, four taxa designated as *S. griseus* (*S. griseus* subsp. *griseus* NRRL B-2307, *S. griseus* subsp. g*riseus* NRRL F-5618, *S. griseus* subsp. *griseus* NRRL F-5621, *S. griseus* subsp. *griseus* NRRL WC-3066) group within the *S. albus* J1074 clade and are thus likely strains of this species and not strains of the streptomycin producer, *S. griseus*.

**Fig. 3.**
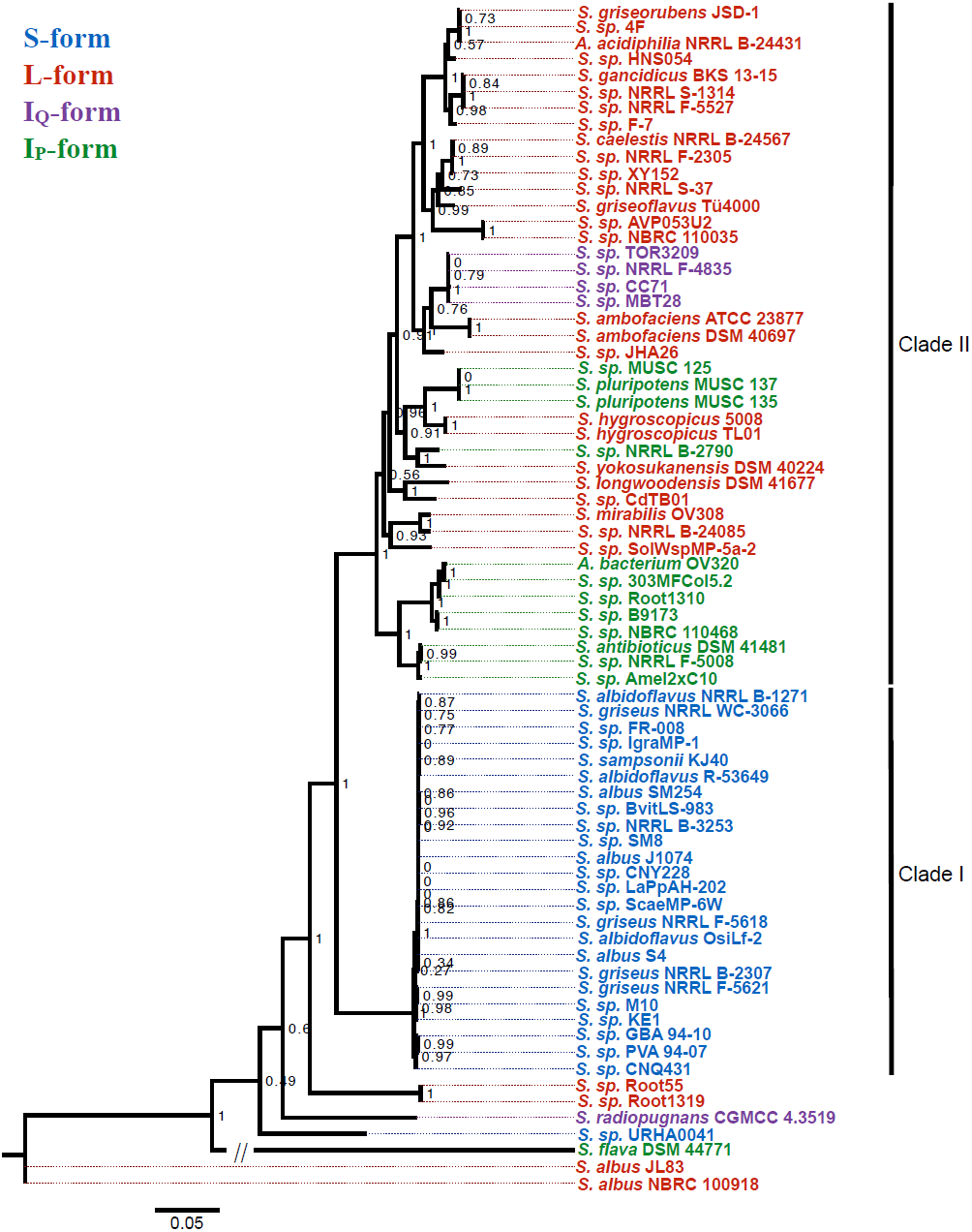
Maximum likelihood phylogeny of 73 *Actinobacteria* analysed in this study. The phylogeny is based on 29 concatenated ribosomal protein DNA sequences. SH-like support values are indicated at nodes as decimal values. Colours indicate gene cluster classification. The scale bar represents 5% sequence divergence.

Next, the phylogenetic tree was colour-coded based on the gene cluster architectures determined above. The sixth bifurcation divides the tree into two major clades, one which contains 24 of the 25 taxa harbouring an S-form antimycin BGC (Clade I), and a second one that contains taxa harbouring exclusively I_P_, I_Q_ and L-form antimycin BGCs (Clade II, Fig. 3). Within Clade I, all 24 S-form taxa are closely related to *S. albus* J1074 and are likely strains of this species. Sequenced strains of *S. albus* are very prevalent within GenBank, which presumably is a consequence of the ease with which they are isolated and cultivated in the laboratory and their ability to strongly inhibit the growth of pathogenic yeasts in bioactivity assays due to antimycin production [25]. Within Clade II, 50% of the L-form antimycin BGCs are harboured by taxa that comprise a single subclade near the top of the tree which includes several isolates from the United States Department of Agriculture NRRL collection as well as, *A. acidiphila* NRRL B-24431 (Fig. 3). The remaining L-form antimycin BGCs are harboured by small groupings of taxa as well as singletons (Fig. 3). Interspersed amongst L-form taxa are those that harbour I_Q_ and I_P_form gene clusters. For instance, four of the five taxa that harbour an I_Q_form antimycin BGC comprise a monophyletic clade situated on a branch sister to that of *S. ambofaciens* ATCC 23877 (Fig. 3), and 11 out of 13 I_P_form antimycin BGCs group into two subclades. Overall, these data suggest that speciation is likely the primary driver for dissemination of the antimycin BGC, which is consistent with the low sequence divergence for taxa closely related to *S. albus* J1074 within Clade I and the formation of gene cluster architecture subclades in Clade II. Interestingly, seven taxa fall outside of Clades I and II: *S.* sp. URHA0041 (S-form), *S. radiopugnans* CGMCC 4.3519 (I_Q_form), *S. flava* DSM 44771 (I_P_ form), and *S.* sp. Root1319, *S.* sp. Root55, *S. albus* JL83 and *S. albus* NRBC 100918 (all L-form), which suggests that these strains may be closely related to the ancestral antimycin producer or that the genes for antimycin biosynthesis were horizontally acquired by these strains (Fig. 3).

### Antimycin BGC phylogeny

A ML phylogeny was inferred from concatenated sequences of *antFGHIJKLMNO* (alignment length 9,736 nt) and colour-coded based on gene cluster architecture as above in order to evaluate evolutionary relationships of antimycin BGCs. These genes were selected because they are conserved in all BGCs and their DNA sequence is not influenced by potential differences in the biochemical diversity their respective biosynthetic pathways may produce. The third bifurcation divides the tree into two major clades, Clade I which harbours only S-form antimycin BGCs and Clade II which harbours exclusively L-I_Q_ and I_P_form antimycin BGCs (Fig. 4). As with the phylosift phylogeny above, 24 of the 25 S-form antimycin BGCs comprise a closely related clade, which was anticipated after the revelation that all of these BGCs are harboured by *S. albus* strains (Fig. 4). L-form antimycin BGCs appear throughout Clade II; 14 of the 30 L-forms clade together at the top of the tree and the majority of the remainder comprise smaller groupings consisting of five, three and two members with two singletons. Four of the five I_Q_form BGCs clade together and are flanked on either side by L-form antimycin BGCs. I_P_form antimycin BGCs form two clades in the center of the tree comprising a total of 12 of the 13 gene clusters. Overall, the tree highlights that phylogenetic placement of antimycin BGCs is linked to their gene cluster architecture in the majority of cases. There are three notable exceptions to this, *S. albus* JL83, *S. albus* NBRC 100918 and *S. sp.* URHA0041 do not group into either Clade I or II and their basal position within the phylogeny may suggest a close relationship with the ancestral antimycin BGC (Fig. 4).

**Fig. 4.**
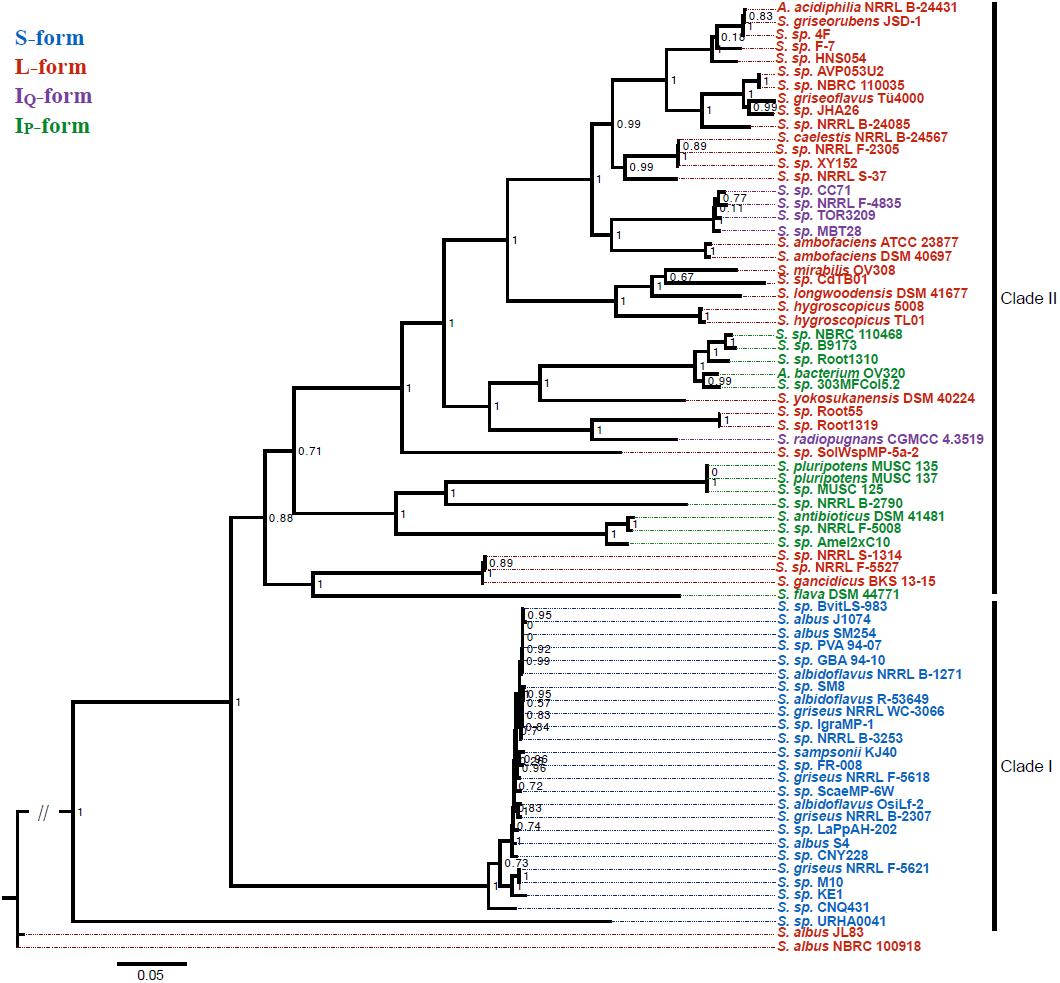
Maximum likelihood phylogeny of the 73 antimycin biosynthetic gene clusters analysed in this study. The phylogeny is based on concatenated *antFGHIJKLMNO* DNA sequences. SH-like support values are indicated at nodes as decimal values. Colours indicate gene cluster classification. The scale bar represents 5% sequence divergence.

### Antimycin BGC evolution

There are obvious similarities between the species and BGC trees. For instance, both trees bifurcate to separate S-form from I_Q_, I_P_ and L-form taxa and BGCs. This combined with the presence of gene cluster architecture subclades in both trees suggests that speciation has been the primary driver for dissemination of the antimycin BGC. With respect to the both the species and BGC trees, it is reasonable to propose that antimycin biosynthesis evolved once and that the ancestral antimycin producer harboured an L-form BGC. To test this hypothesis, a likelihood analysis was used to predict the ancestral node for each architecture of the antimycin BGC based on its distribution within the phylosift phylogeny. As expected, the likelihood analysis predicted that the ancestral antimycin producer harboured an L-form BGC (Fig. 5). This supports a model in which loss of *antP* and/or *antQ*, rather than their frequent independent acquisition, resulted in diversification of gene cluster architecture. The ancestral antimycin producer likely gave rise to both *S. albus* JL83, *S. albus* NBRC 100918 and *S. sp.* URHA0041, however *S. sp.* URHA0041 lost *antP* and *antQ* after speciation. The same ancestral L-form strain presumably also seeded Clade I where *antP* and *antQ* were lost in the process, but were retained during the genesis of Clade II. One major diversification event likely occurred to give rise to most of the I_P_form antimycin BGCs, however a second diversification event appears to have occurred where *antQ* was lost by the ancestor of *S. sp.* NRRL B-2790, *S. pluripotens* MUSC 135, *S. pluripotens* MUSC137 and *S. sp.* MUSC125, but was retained by the two *S. hygroscopicus* strains that the aforementioned clade with. Finally, four of the five I_Q_form BGCs are derived from the L-form ancestor of *S. ambofaciens* ATCC 23877 and *S. ambofaciens* DSM 40697.

**Fig. 5.**
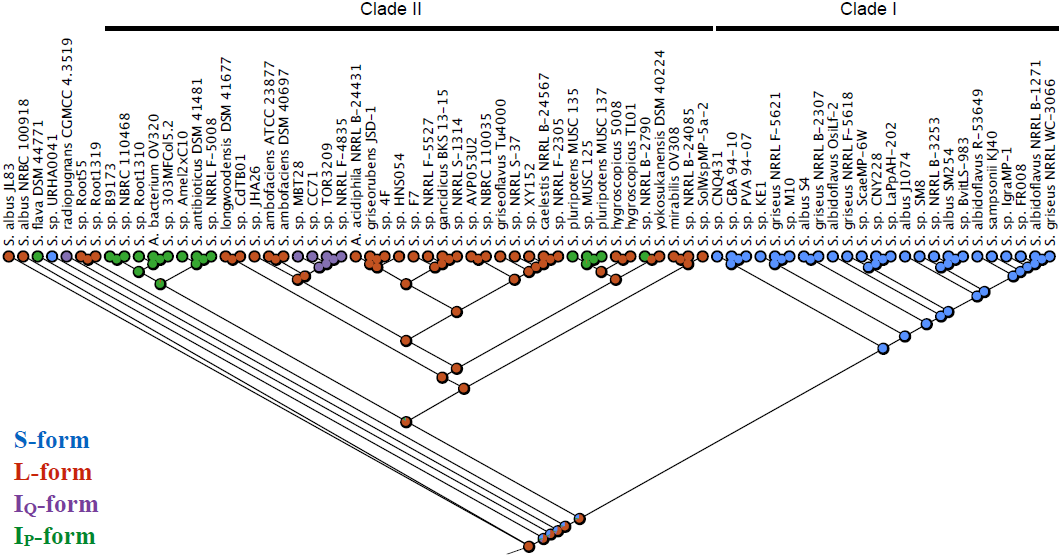
Likelihood analysis of ancestral antimycin BGC architectures. Filled circles are colour-coded to represent the proportional likelihood of BGC architecture at ancestral nodes.

Although the phylosift and BGC trees are consistent with that proposed above, there is noticeable discordance between the two phylogenies. The most profound of these are highlighted below. Three closely related taxa, *S. gancidicus, S. sp.* NRRL S-1314 and *S. sp.* NRRL F-5527 group within the large subclade of L-form BGCs in Clade II of the phylosift tree, but shift to occupy a distantly related lineage in Clade II of the *antFGHIJKLMNO* phylogeny. This suggests that the parent of the *S. gancidicus* subclade likely received its L-form gene cluster by horizontal gene transfer. The same is also likely true for *S. sp.* NRRL B-24085, which is located within the centre of Clade II in the phylosift tree, but then joins the large L-form subclade of Clade II in the BGC tree. Interestingly, *Sacharropolyspora flava* DSM 44771, which is an ‘outlier’ in the phylosift tree (i.e. does not group within Clade I or II) becomes part of Clade II in the BGC tree and shares an ancestral node with the three-membered *S. gancidicus* subclade described above. This suggests that *S. flava* and the *S. gancidicus* subclade likely received their antimycin BGC from the same or a closely related ancestor and is consistent with the hypothesis that *S. flava* originally harboured an L-form BGC, but independently lost *antQ* to give rise to its I_P_form BGC. In addition, three other outliers from the phylosift tree group within Clade II of the BGC tree: *S. sp.* Root1319, *S. sp.* Root55 and *S. radiopugnans*, which suggests that their antimycin BGC was horizontally acquired and moreover that *S. radiopugnans* likely independently evolved an I_Q_form antimycin BGC from the clade founder.

### Conclusions and perspectives

In this study, 73 antimycin BGCs were identified in the genome sequences of *Actinobacteria* deposited in GenBank. The isolation data for these strains indicates that antimycin-producing actinomycetes are likely globally distributed, highlighting an important role for inhibiting cytochrome c reductase in diverse ecological niches. The majority of the antimycin BGCs identified contained both the *antP* kynureninase and the *antQ* PPTase (L-form) or neither of these (S-form), while a minority of the gene clusters lacked either the *antP* or *antQ* (I_Q_ or I_P_form, respectively). Phylogenetic analyses revealed two distinct lineages separating S-form from L-, I_Q_ and I_P_form taxa and BGCs and although a handful of taxa appear to have acquired the antimycin BGC via horizontal gene transfer, the primary means for dissemination of the gene cluster is vertical transmission. The contextual requirement of *antP* and *antQ* presumably permitted divergent evolution of the antimycin biosynthetic pathway. We propose that the ancestral antimycin producer harboured an L-form antimycin BGC, which spawned two main clades, one composed of S-forms that lack both *antP* and *antQ*, and one composed of L-forms with distinct subclades of I_P_ and I_Q_forms (Fig. 6).

**Fig. 6.**
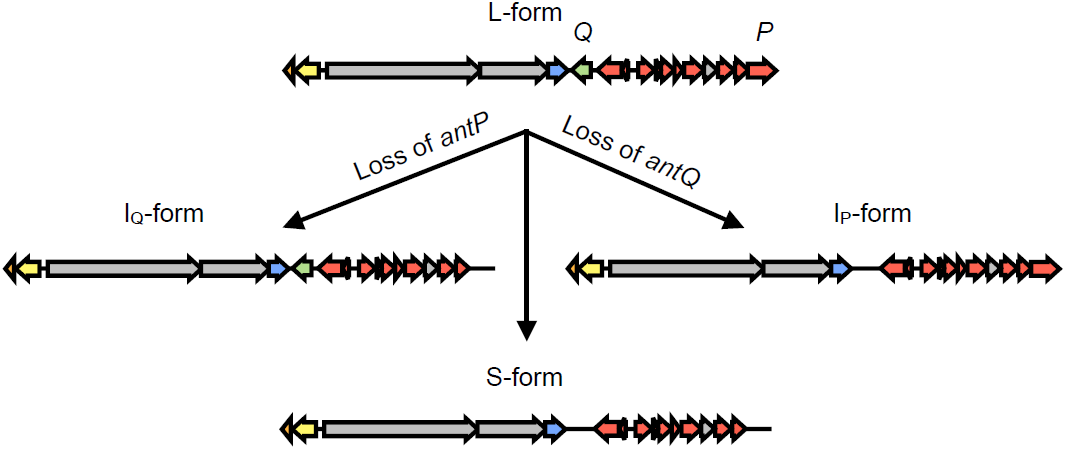
Proposed evolutionary path of the antimycin BGC. The ancestral antimycin producer likely harboured an L-form gene cluster which independently lost either *antP, antQ* or both of these genes to give rise to the I_Q_, I_P_ and S-form gene clusters, respectively.

### Materials and methods

### Identification of putative antimycin biosynthetic gene clusters

The genomes available in GenBank on the 9^th^ of May 2017 for select genera of *Actinobacteria* (*Actinobactera, Actinomadura, Actinospica, Amycolatopsis, Kitasatospora, Micromonospora, Nocardia, Saccharopolyspora, Planomonospora, Pseudonocardia, Salinispora, Streptacidiphilus,* and *Streptomyces*) were downloaded using the ncbi-genome-download python script provided by Kai Blin available at <u>https://github.com/kblin/ncbi</u>-<u>genome-download</u>. One thousand four hundred and twenty-one genomes were downloaded in total and were subsequently annotated using Prokka 1.12 [18]. The Unix command grep was used to remove any line containing the string “/gene” and then the Unix command sed was used to edit “/locus_tag” to “/gene”. A multigeneblast database was created using the makedb programme of multigeneblast version 1.1.13 [19]. The genes *antFGHIJKLMNOPQ* of the antimycin BGC from *S. ambofaciens* ATCC 23877 [16] were used as a multigeneblast query with the default settings. The resulting output was inspected manually to identify genomes harbouring a putative antimycin BGC. PROmer [26] and the *S. ambofaciens* antimycin BGC were used to identify contigs comprising antimycin BGCs split across more than one contig. This applied to the following taxa: *S. gancidicus* BKS 13-15, *S. sp.* B9173, *S. sp.* CC71, *S. sp.* HNS054, *S. sp.* IgraMP-1, *S. sp.* MBT28, *S. sp.* NRRL B-24085, *S. sp.* TOR3209, *S. sp.* SM8. *S. wadayamensis* strain A23, which harbours a putative S-form antimycin BGC, was discarded because it lacked multiple phylosift markers (see below).

### Phylogenetic analyses

In order to infer a species phylogeny, 29 single-copy phylogenetic markers (13,061 nt in total) were identified and extracted using Phylosift version 1.0.1 [Table S1, [27]] and concatenated in the order: DNGNGWU00002, DNGNGWU00003, DNGNGWU00007, DNGNGWU00009, DNGNGWU00010, DNGNGWU00011, DNGNGWU00012, DNGNGWU00014, DNGNGWU00015, DNGNGWU00016, DNGNGWU00017, DNGNGWU00018, DNGNGWU00019, DNGNGWU00021, DNGNGWU00022, DNGNGWU00023, DNGNGWU00024, DNGNGWU00025, DNGNGWU00026, DNGNGWU00027, DNGNGWU00028, DNGNGWU00029, DNGNGWU00030, DNGNGWU00031, DNGNGWU00033, DNGNGWU00034, DNGNGWU00036, DNGNGWU00037, DNGNGWU00040. The concatenated phylogenetic marker sequences were aligned using 16 iterations of MUSCLE [28] and the. fasta format alignment was converted to sequential phylip format using Geneious R8.1.19. Phylogenetic relationships were inferred from this alignment using the web-implementation of PhyML3.0 available at <u>http://www.atgc-montpellier.fr/phyml/</u> [29].

In order to infer a phylogeny for putative antimycin BGCs, 10 genes (*antFGHIJKLMNO*) were extracted and aligned using MUSCLE. The resulting alignment was imported into Geneious R8.1.19 and manually trimmed to the same length prior to concatenating sequences in the following gene order: *antFGHIJKLMNO.* The concatenated alignment was then converted to sequential phylip format and a phylogenetic tree was inferred using PhyML3.0 as above.

### Likelihood analysis

Reconstruction of the ancestral state was performed essentially as described previously [22]. Briefly, the trace character function of Mesquite v3.2 [30] was used to infer the ancestral node for the antimycin BGC within the species tree. A categorical character matrix for BGC type was created and likelihood calculations were performed using the Mk1 model.

## Acknowledgments

We thank Bohdan Bilyk, Divya Thankachan and Liam Sharkey for their helpful discussions during this study. This work was in part undertaken on ARC2 and MARC1, part of the High Performance Computing facilities at the University of Leeds. This work was funded by a grant from the Biotechnology Sciences Research Council grant (BB/N007980/1) awarded to RFS. The funders had no role in study design, data collection and interpretation, or the decision to submit the work for publication.

